# Linkage Analysis in Caribbean Hispanic Families with Puerto Rican Ancestry Idenitfies an Alzheimer Disease Locus on chromosome 9

**DOI:** 10.1101/2020.03.10.986083

**Authors:** Farid Rajabli, Briseida E. Feliciano-Astacio, Holly N. Cukier, Liyong Wang, Anthony Griswold, Kara L. Hamilton-Nelson, Larry D. Adams, Vanessa C. Rodriguez, Pedro Ramon Mena, Sergio Tejada, Katrina Celis, Patrice L. Whitehead, Derek J Van Booven, Natalia K. Hofmann, Parker Bussies, Michael Prough, Angel Chinea, Nereida I Feliciano, Heriberto Acosta, Clifton L. Dalgard, Jeffery M. Vance, Michael C. Cuccaro, Gary W. Beecham, Margaret A. Pericak-Vance

## Abstract

**Background:** The ancestral genetic heterogeneity (admixture) of Caribbean Hispanics makes studies of this population critical to the discovery of ancestry-specific genetic factors in Alzheimer disease. In this study, we performed whole genome sequencing in multiplex Caribbean Hispanic Puerto Rican families to identify rare causal variants influencing Alzheimer disease through linkage and segregation-based approaches.

**Methods:** As part of the Puerto Rican Alzheimer Disease Initiative, whole genome sequencing data were generated for 100 individuals (61 affected) from 23 Puerto Rican families. To identify the genetic loci likely to carry risk variants, we performed a parametric multipoint affected individuals-only linkage analysis using MERLIN software. Following the linkage analysis, we identified the consensus region (heterogeneity logarithm of the odds score (HLOD) > 5.1), annotated variants using Ensembl Variant Effect Predictor, and combined annotation dependent depletion score (CADD). Finally, we prioritized variants according to allele frequency (< 0.01), function (CADD > 10), and complete segregation among affected individuals.

**Results:** A locus at 9p21 produced a linkage HLOD score of 5.1 in the parametric affecteds-only multipoint affected individuals-only model supported by 9 families. Through the prioritization step, we selected 36 variants (22 genic variants). Candidate genes in the regions include *C9orf72, UNC13B*, and *ELAVL2*.

**Conclusions:** Linkage analysis of Caribbean Hispanics Puerto Rican families confirmed previously reported linkage to *9p21* in non-Hispanic White and Israeli-Arap families. Our results suggest several candidates in the region as conferring AD risk. Identified putative damaging rare variants in multiplex families indicates the critical role of rare variation in Alzheimer disease etiology.

## 1. Introduction

Alzheimer disease (AD) is the leading cause of dementia in the elderly and occurs among all ethnic and racial groups^1^. Genetic studies across populations have shown different risk effect size at the same loci and different risk loci^2,3^. The *APOE* and *ABCA7* genes are notable examples illustrating this population-specific variability. Association studies show that the effect of the *APOE* ε4 risk allele in East-Asian and non-Hispanic white (NHW) populations is strong, while it is considerably lower in populations with African ancestry^2^. Reitz et al. found that the *ABCA7* gene has a much stronger risk in African Americans (AA) than was found in NHW individuals^4^. In a follow-up study, Cukier and colleagues identified a 44 base pair (bp) deletion in *ABCA7* associated with AD specific to individuals with an African ancestral background^5^. These findings show that association between risk and protective factors for AD may be different across populations and suggest that distinct genetic architecture among race/ethnic groups may underlie disease pathogenesis. Thus, studying diverse populations may yield new insights into genetic mechanism of AD pathology that are essential to understanding AD genomics and identifying targets for the effective treatment. However, the majority of genetic studies in AD to date have focused NHW populations^6^.

Additionally, currently identified AD loci only account for a portion of the underlying genetic basis of disease. One of the explanations to account for the missing heritability is the rare variant hypothesis; that is, rare genetic variation substantially contributes to disease risk^7^. One possible solution to identify rare risk variants is to use family-based whole genome sequencing (WGS) approach^8^. Multiplex families (families with several individuals affected with AD) have a higher genetic burden of disease than sporadic cases. Furthermore, using multiplex families allows for the identification of highly penetrant rare variants that are shared among several affected individuals in a family. Thus, sequencing multiplex families provide a powerful strategy to identify rare risk variants associated with AD.

To begin addressing these issues, we focused on a <u>C</u>aribbean <u>Hi</u>spanic <u>P</u>uerto <u>R</u>ican (CHI PR) population and selected multiplex AD families. CHI PRs have a high prevalence rate of AD (12.5%) that is higher than it was estimated in the general U.S. population (10.1%), but CHI PRs are underrepresented in AD genetic studies^9^. Moreover, the genetic make-up of CHI PRs is not homogenous; they are highly admixed, with average ancestry values of 69% European (EU), 17% African (AF), and 14% Native American (AI)^9^. The ancestral heterogeneity in CHI PR population make it unique in the discovery of novel AD loci, as well as in the replication of findings from European-descent and African-descent populations. In this study, we performed WGS in multiplex CHI PR families to identify highly penetrant rare variants influencing AD through linkage and segregation-based approaches.

## 2. Methods

### 2.1 Study Samples

Study participants were recruited as part of the Puerto Rico Alzheimer Disease Initiative (PRADI). Families have been ascertained through the John P. Hussman Institute for Human Genomics at the University of Miami. Families were required to have multiple (at least 2) members with AD, available genomic DNA, and *APOE* genotypes. Patients were evaluated to determine their AD clinical status based on the National Institute of Neurological and Communicative Disorders and Stroke-Alzheimer’s Disease and Related Disorders Association criteria^10,11^. A detailed description of our ascertainment and clinical evaluation methods of the individuals have been previously described^9^. However briefly, cases were defined as individuals have a previous clinical diagnosis of AD, mild cognitive impairment (MCI), dementia, or show evidence of a memory disorder, and meet standard criteria for AD or MCI^10-12^. We excluded individuals whose memory and cognitive problems are secondary to other causes (e.g., stroke, psychoses, etc.) and those with a known mutation (e.g., *APP*^13,14^, *PS1*^15-18^, or *PS2*^19,20^). To be included as an unaffected family members had to meet the same inclusion criteria as the controls in addition to being a first- or second-degree relative of a case. An adjudication panel consisting of a neurologist, neuropsychologist and clinical team members reviewed all medical records, clinical and family history and clinical tests. The panel reviewed all data and assigned best-estimate diagnoses. To be classified as AD individuals had to meet the current NIA-AA criteria^12^. They were further classified as definite (neuropathologic confirmation), probable, or possible AD. Diagnoses of MCI were assigned using the NIA-AA criteria^10^. Cognitively normal individuals with no history of memory problems and MMSE or 3MS scores that fall above clinical cutoffs were designated as unaffected at the current age at exam.

Twenty-three CHI PR families (100 total individuals, 61 affected individuals) were selected to perform the initial linkage analysis (Table 1). This study was carried out in accordance to the recommendations of the National Institute of Health Guiding Principles for Ethical Research Pursuing Potential Research Participants Protection and the 2016 National Institute of Health Single Review Board (sIRB) Policy. This study received ethical approval from University of Miami Institutional Review Board (approved protocol #20070307) and Universidad Central del Caribe Institutional Review Board (approved protocol # 2016-26). The Universidad Central del Caribe is relying on the designated UM-IRB by an Institutional Review Board Authorization Agreement (Protocol: Genetic Studies in Dementia). All subjects (participant or proxy) gave written informed consent. This study was carried out in accordance with the Declaration of Helsinki and amendments.

**Table 1.** Puerto Rican family sample characteristics.

| **Characteristic** |  |
| --- | --- |
| Number of families, | 23 |
| Subjects in families, | 100 |
| Affection Status, (%) |  |
| Affected | 61 (61%) |
| Unaffected | 39 (39%) |
| Proportion of Women, (%) | 77 (77%) |
| Mean age of onset of affecteds, years (SD) | 73.9 (11.5) |
| Mean age at last examination of unaffecteds, years (SD) | 68.8 (9.8) |

### 2.2 Array genotyping, whole genome sequencing, and annotation

Genome-wide genotyping data were obtained using two platforms: the Illumina Multi-Ethnic Genotyping Array and the Illumina Global Screening Array. Quality control (QC) analyses were performed using software PLINK v.2^21^. Variants with the call score less than 95%, or not in Hardy-Weinberg equilibrium (HWE) (p<1.e-6) were eliminated. The individuals with the genotyping call rate less than 97% were removed. The concordance between reported sex and genotype-inferred sex was checked using X-chromosome data. The relatedness among the individuals were assessed using “identical by descent” (IBD) allele sharing.

WGS was generated at the Uniformed Services University of the Health Sciences (USUHS) and Center for Computational Sciences at the University of Miami using a Illumina NovaSeq instrument (Illumina, San Francisco, California, USA). Raw FASTQ files were processed on a high-performance computing cluster maintained by the Center for Computational Sciences at the University of Miami. We performed Genome Analysis Toolkit (GATK) best practices on the data. For the quality control of the FASTQ files, we used FASTQC software^22-24^. Reads were aligned to the GRCh38 using bwa-mem, duplicates marked using PICARD, and base qualities recalibrated. Multi-sample variant calling and joint genotyping were performed using the GATK HaploteCaller across all samples from the study^25,26^.

Variants were annotated for their genomic location, relationship to genes, putative function, combined annotation dependent depletion (CADD) score, and allele frequency using in-house pipeline based on Variant Effect Predictor Ensembl framework and Bravo^27-29^.

### 2.3 Linkage analyses

We performed genome-wide multipoint parametric and nonparametric linkage scans on the array data using MERLIN v 1.1.2 software^30^. First, we pruned markers using the PLINK software v2^21^ with independent pairwise LD pruning option (r^2^ < 0.02). Then, we extracted allele frequencies for all markers using PR dataset (PUR) in the 1000Genome data^31^. Next, we performed both parametric multipoint affected individuals-only and nonparametric linkage analyses. Parametric multipoint affected individuals-only analysis was performed assuming dominant model with incomplete penetrance (non-carrier 0.01, and carriers 0.90) and rare disease allele frequency 0.0001. Nonparametric linkage analyses was carried by using NPL-pair and NPL-all statistics from Whittemore and Halpern, and Kong and Cox linear model of logarithm of the odds (LOD) scores^32,33^. The significance linkage threshold of LOD > 3.3 (p-value = 4.9e-5) was defined for the consensus regions (with the multiple families contributing to the linkage peak) as it was recommended by the Lander and Kurglyak^34^.

Significant linkage regions were further used to examine the identical by descent (IBD) regions shared across all AD individuals within the families contributing to the linkage peak (LOD > 0.57). IBD regions were estimated using MERLIN software and illustration was performed using PROGENY software^35^.

### 2.4 Variant filtering and prioritization

For the variants located within the linkage region, we applied several filtering steps to prioritize them. Firstly, we selected variants that showed complete segregation among the affected individuals within the families. Secondly, we filtered variants with the minor allele frequency (MAF) less than 0.01 using TopMed allele frequency information. Then, we used a CADD score of greater than 10 to select the top 10% of deleterious variants in the human genome. Finally, we prioritized the variants by selecting variants based on their presence in multiple families, or multiple variants falling within the same gene, or both.

### 2.5 Genotyping validation

We performed Sanger sequencing^36^ to validate the prioritized variants and to examine previously reported repeat expansion at the region linked to disease^37,38^. The details of the Sanger sequencing protocol are in appendix 1.

## 3. Results

To recognize rare risk variants that segregate with AD and potentially identify novel candidate loci in CHI PR population, we analyzed WGS data in multiplex CHI PR families. We performed two-step approach: linkage analysis to find the chromosome regions likely to harbor rare risk variants with a large effect size and then a segregation-based approach to identify rare causal variants.

### 3.1 Linkage results

The parametric multipoint linkage scan identified a significant positive linkage signal with the maximum HLOD = 5.1 on chromosome 9p21. In total, nine families supported evidence of linkage to the linkage region on 9p21 with LOD scores higher than or equal to 0.57 (corresponding to a nominal *P* value of 0.05). The 1-LOD unit down region (approximately 95% confidence interval of the linkage region) spans between positions 32 Megabases (Mb) and 38 (Mb) on chromosome 9^39^. The IBD sharing analysis among the families supporting the linkage findings defined region of interest that includes chromosomal region from 23 Mbto 39 Mb (Figure 1). Non-parametric linkage analysis showed the strongest signal in the same region (9p21) with the LOD score 2.3.

**Figure 1.**
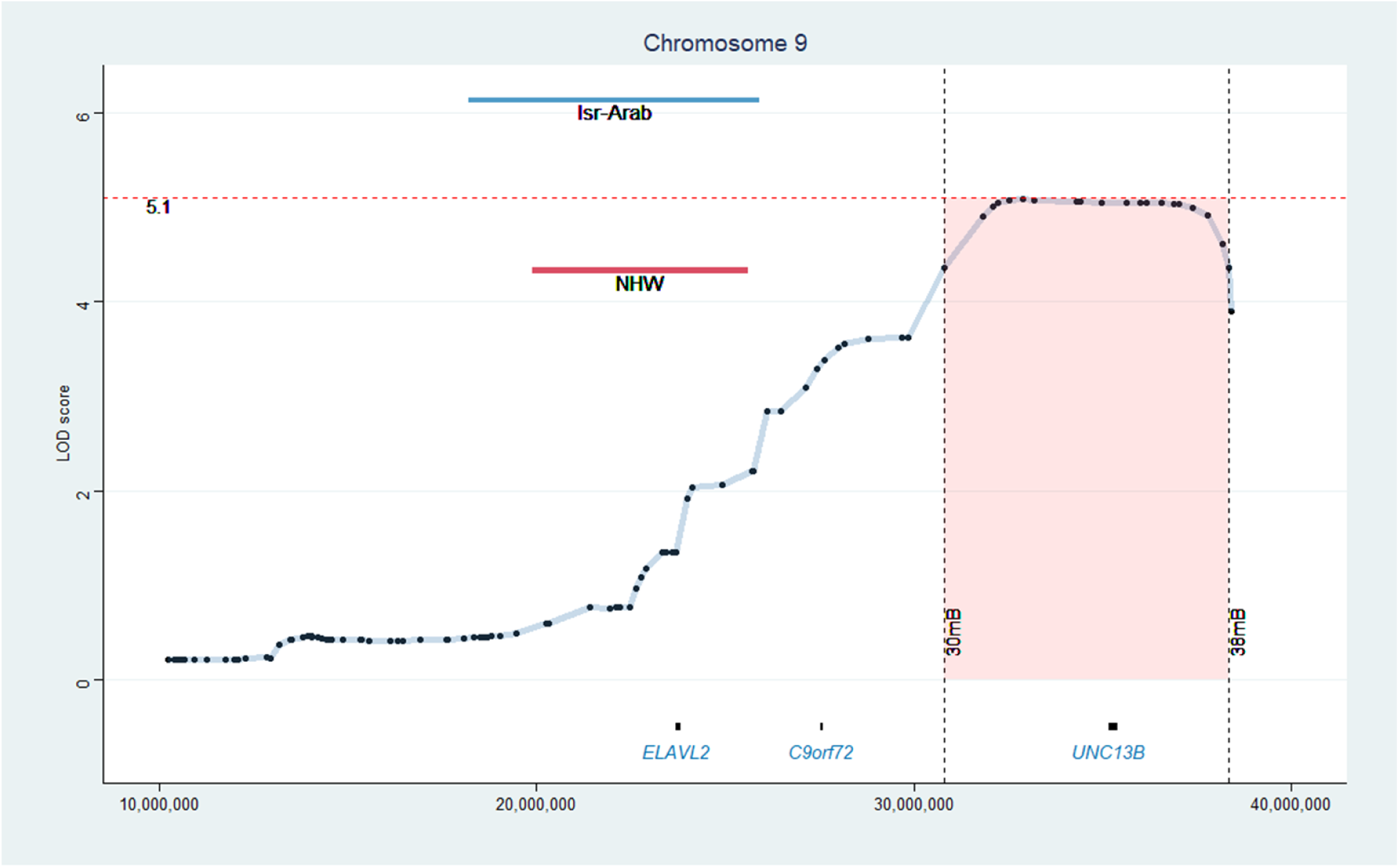
Multipoint linkage peak on chromosome 9. x-axis denotes physical position in base pairs, and y-axis denotes the LOD score. Blue and red lines illustrate the linkage results from independent Israeli-Arab and non-Hispanic Whites family studies. Lines on the x-axis align with the linkage region and on the y-axis align with corresponding LOD score.

### 3.2 Prioritization results

We identified 116 variants that segregated with disease in affected individuals in at least 1 of the 6 families that contributed to the linkage peak, with the rare allele frequency (< 0.01), and moderate-to-high CADD score (>10). Out of 116 variants, 45 (38.8%) were intergenic and 71 (61.2%) were genic. We further prioritized our findings using additional filtering steps such as the absence of the variants in unaffected individuals in the family, the same variant being identified in multiple families or multiple variants occurring in the same gene, or both. We ended with the 36 “high priority” variants (22 genic variants). Table 2 summarize the prioritized 22 genic variants and classifies them according to genes and provides average CADD score and number of families they have been segregated. Top genes include at least 2 high priority variants and segregated at least in two families. Remarkably, six of the 22 genic variants were identified in the gene *UNC13B* that segregate with AD in four families with the average CADD score 16.2. *UNC13B* was the only gene with multiple, segregating coding changes: rs35199210 (Asp238Glu) and rs41276043 (Phe1096Leu). *In silico* evaluation found CADD scores for both > 22, and Polyphen predicted both to be highly deleterious. The rs35199210 (Asp238Glu) segregates with the disease in a family with 6 AD individuals. The full list of the prioritized variants is included in the supplement material.

**Table 2.**
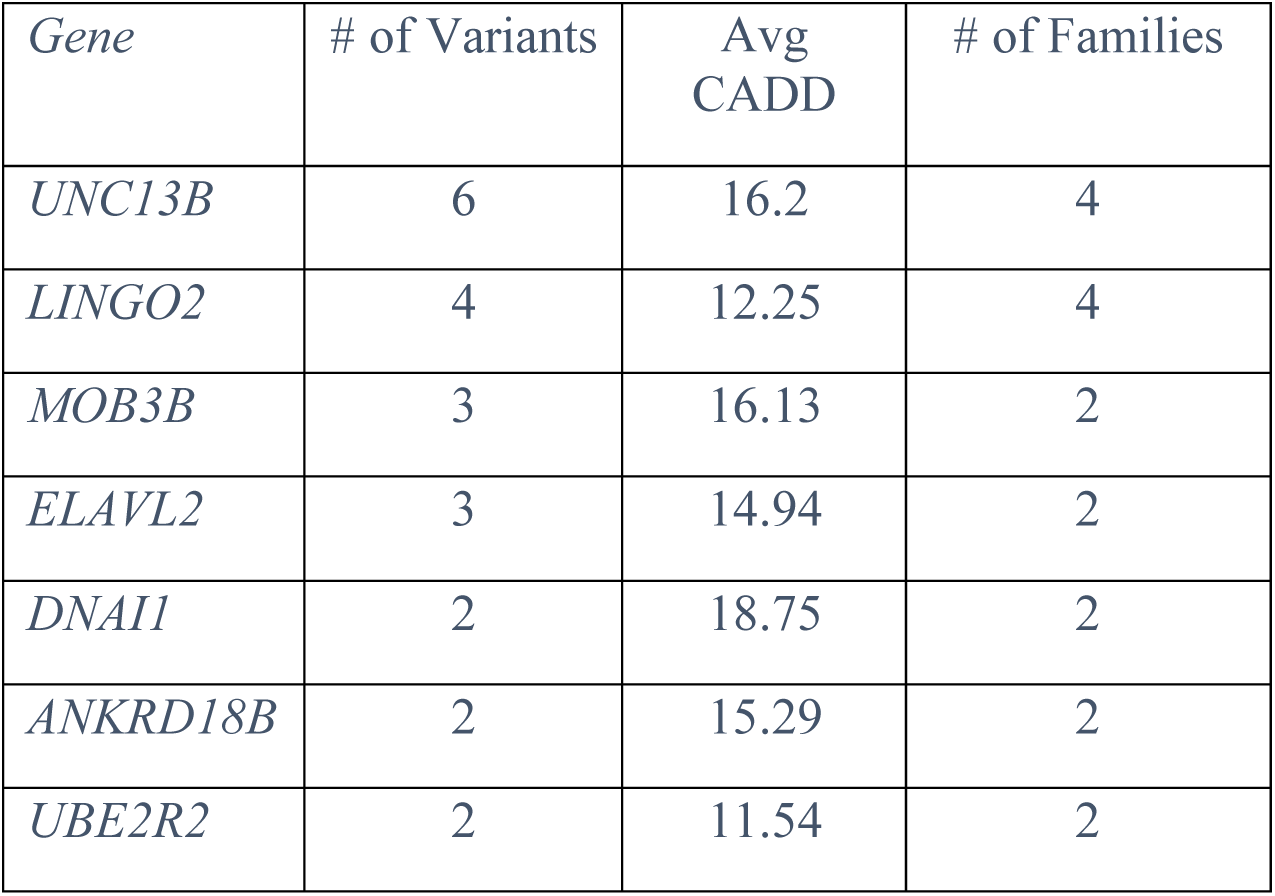
Prioritized variants from the linkage region.

### 3.3 Validation results

We validated all prioritized variants by Sanger sequencing. Additionally, we evaluated whether any of the families carried the hexanucleotide repeat expansion in *C9orf72* gene^40^. The hexanucleotide repeat in *C9orf72* lies within the 9p21 linkage region, and known as the most important genetic cause of familial frontotemporal dementia and amyotrophic lateral sclerosis^37,38^. Results showed that no family had extended repeat expansions (>30 repeats).

## 4. Discussion

We report a genome-wide significant multipoint linkage region (HLOD > 5.1) identified by analysis of 23 CHI PR multiplex families. The region of interest overlaps with the previously reported AD linkage peaks in the independent set of NHW^41^ and Israeli-Arab families^42^ (Figure 1). Additionally, the 9p21 region was also linked to other neurodegenerative diseases such as frontotemporal dementia and amyotrophic lateral sclerosis^37,38^. A segregation-based approach in this study identified numerous rare, predicted damaging rare risk variants. There are two interesting genes that came out from the high priority variants list (Table 2): *unc-13 homolog B* (*UNC13B and ELAV like RNA binding protein 2* (*ELAVL2*).

*UNC13B* encodes for a protein involved in calcium regulated vesicle priming and release at excitatory/glutamatergic synapses^43^. Variants in the *UNC13B* gene have also been reported in individuals with bipolar disorder and schizophrenia, while another member of the gene family, *UNC13A*, has been implicated in amyotrophic lateral sclerosis^44-47^. There are two missense variants in the gene *UNC13B* that segregate with AD in families, suggesting that *UNC13B* may be a candidate AD risk gene. Functional *in silico* analysis predicted these variants to be among the top 1% most deleterious variations.

*ELAVL2* is a neural-specific RNA binding protein coding gene that regulates the alternative splicing process and has been previously related to neurodegenerative diseases, including AD^48^. Neuronal members of the *ELAVL* protein family have been shown to play an important role in the regulation of neuron-specific alternative splicing of *APP* pre-mRNA^49^. The alteration of the post-transcriptional *APP* expression is critical in the pathogenesis of AD.

This study has potential limitations. First, the accuracy of the structural variants calling is not high. Although we sequenced the known repeat within the *C9orf72* gene, other potential structural variants need to be examined. A different copy number of variations in *C9orf 72* may contribute to the AD linkage signal. Second, we used an *in silico* tool to predict the pathogenicity of the variants. Even though this approach is useful to prioritize the putative variants, *in vitro* functional studies on characterization of these variants needed to be undertaken.

The identification of putative damaging rare variants in multiplex families indicate the critical role of rare variation in AD etiology. Linkage analysis of CHI PR families confirmed previously reported linkage to 9p21 in NHW and Israeli-Arab families. Our results did not support the known candidate locus *C9orf72* gene as the AD gene contributing to risk in our families. The data did suggest the presence of AD risk variants in the *UNC13B* and *ELAVL2* genes. Further directions include a comprehensive structural variants analysis and *in vitro* characterization of the prioritized variants.

## Acknowledgements

We thank the patients and families that participated in the study. This investigation was supported by grant AG054074 and AG057659 from the National Institutes on Aging of National Institutes of Health.

